# Local people’s perceptions on the influence of linear infrastructure on human-wildlife conflict in Southern Kenya

**DOI:** 10.1101/2025.08.05.667126

**Authors:** Lucy Waruingi, Alistair Rob Marchant, Diego Juffe, D. Neil Burgess, O. Tobias Nyumba, Henry Gandhi Odhiambo

## Abstract

Globally, conservation and development efforts are impacted by human-wildlife conflict as this issue negatively affects human lives. Increasingly, sub-Saharan Africa has become the focus of linear transportation infrastructure development that encroaches on many traditional wildlife areas. One of the resulting effects of infrastructure development – allowing people to move into areas they were not previously living – can be human-wildlife conflict. This relationship between people, wildlife and infrastructure has not been well investigated in Kenya. We studied local people’s perceptions of the impacts of the Mombasa to Nairobi Standard Gauge Railway (SGR), roads, oil pipelines and electricity transport lines on human-wildlife conflict in southern Kenya. Our results show that people’s perceptions are that human wildlife conflict has increased in the past five years due to the construction of the SGR, but not in relation to other infrastructure. We have no independent data to check this result, and there is significant politics around this railway project, but the result is robust along 100s of kilometres of railway construction route. Our conclusion is that addressing human-wildlife conflict requires a holistic overview and understanding of the drivers and their interactions to support the development of strategies that support human and wildlife coexistence.

## 1. Introduction

Human-wildlife conflict (HWC) is one of the most intractable conservation and development challenges (Madden 2004, Shrestha *et al*. 2007, Lamarque *et al*. 2009, Nyumba *et al*. 2020). HWC manifests in different direct and indirect forms including loss of human life, damage to crops and property, livestock depredation and retaliatory killing of wildlife leading to loss of livelihoods, psychological and economic well-being, and food security for people and loss of biodiversity and changes in ecosystem structure (Nyumba *et al*. 2020). HWC is extremely widespread in ecosystems characterised by high ecological and anthropogenic drivers such as wildlife movement patterns, land-use change and infrastructure development acting singly or in combination resulting in a complex web of interactions (Lamarque *et al*. 2009, Hoare 2015a, Nyumba *et al*. 2021).

The past decade has witnessed a surge in the growth of linear transportation infrastructure (LTI) in sub-Saharan Africa, including pipelines, roads and railways (Dulac 2013, Lynam *et al*. 2014,Alamgir *et al*. 2017, DCP Kenya 2019). Some of this can be traced to the implementation of the Chinese government’s global ‘Belt and Road’ policy (Kaplinsky *et al*. 2007). While LTI can promote social and economic development, it can also deliver negative outcomes for wildlife and people (Venter *et al*. 2016, Mahmoud *et al*. 2017, Okita-Ouma, Lala, Moller, *et al*. 2017, Teo *et al*. 2019). This is particularly the case when LTI enters areas rich in wildlife and promotes human movements into these areas (Ree Rodney *et al*. 2011, Laurance *et al*. 2015, Nyumba *et al*. 2021). Case studies on HWC highlighting the influence of various ecological and anthropogenic drivers are found throughout the world (Nyhus 2016). The negative effects of LTI on wildlife have been documented for decades and include habitat loss and fragmentation, wildlife mortality, disturbance and creation of barriers to the movement and migration of wildlife, also referred to as the *“barrier effect”* (Ree Rodney *et al*. 2011, Morelle *et al*. 2013, Schwartz *et al*. 2020). One of the consequences of the barrier effect of LTI is wildlife mortality during crossing attempts and behavioural avoidance (Anderson, 2002; Forman et al., 2003). This reduces access to necessary resources and may deflect wildlife into surrounding human settlements leading to incidents of HWC.

In recent years, however, awareness and discussion of the influence of technology, transportation and energy on human-wildlife interactions have increased. These draw on studies around transportation-related collisions also known as wildlife-vehicle collisions (WVCs) (e.g. Coelho *et al*. 2008, Matos *et al*. 2012, Lala *et al*. 2021) and the emerging renewable energy sector investments including oil exploration and exploitation and the installation of energy turbines and industrial solar infrastructure (Nyhus 2016). These studies have not related HWC to the growth and expansion of infrastructure, especially the linear transportation infrastructure in HWC hotspots.

Over the past ten years, Kenya has experienced particularly rapid and extensive infrastructure development under the country’s ‘Vision 2030’ Development agenda. During this period, the country constructed the Standard Gauge Railway (SGR) connecting the coastal town of Mombasa to towns in western Kenya (Nyumba *et al*. 2021). The SGR runs parallel to other linear infrastructure including oil pipelines, power lines and highways and cuts through two important landscapes – Tsavo and Shimba Hills – and associated protected areas. These two landscapes are not only important for wildlife conservation but also support local human populations living within community and private conservancies present along the boundaries of the protected areas. The landscapes are currently facing multiple threats from land-use change, including existing, new and planned infrastructure development, the SGR being the iconic national investment across the landscape.

We aim to characterise HWC and determine whether distance from infrastructure, and in relation to other factors, influences the occurrence of HWC. The discussion focuses on results from a case study along the SGR within the Shimba Hills and Tsavo landscapes in southern Kenya. The research questions are as follows: (1) What are the types and nature of HWC in the Shimba and Tsavo landscapes? (2) To what extent are these conflicts differentially borne by residents of different counties? And (3) what is the role of distance from the target infrastructure, and social and economic drivers on the occurrence of HWC? Answers to these questions will facilitate deeper insights into linear infrastructure design, construction and operation that delivers better biodiversity planning and management, including the resolution and mitigation of HWC in Kenya.

## 2. Methods

### 2.1 Study area

This study builds on the previous work of Nyumba *et al*. (2021) which assessed the ecological impacts of infrastructure along the entire phases I & IIA of the SGR and established through group interviews and meetings that intensification of HWC was one of the consequences of the construction and operation of the SGR and associated infrastructure in Kenya. The present study was conducted within a 20km buffer of the SGR covering a land-use mosaic of wildlife conservation, human settlement and agriculture from Mombasa to Makueni counties (Figure 1). The study area comprises parts of the Shimba Hills and Tsavo protected areas. The Shimba Hills run along the south coast of Kenya in Kwale County and are composed of diverse, threatened and endangered biodiversity within a landscape ranging from forests, bushlands, and grasslands to wetlands. The area includes protected areas such as the Shimba Hills National Reserve, Mkongani Forest and Mwaluganje Conservancy making up a total of 253 km^2^ (Malonza *et al*. 2018). The landscape experiences a humid semi-equatorial climate with an average monthly temperature of between 24°C and 28°C and rainfall of 1200mm per annum with long rains between March to June.

**Figure 1:**
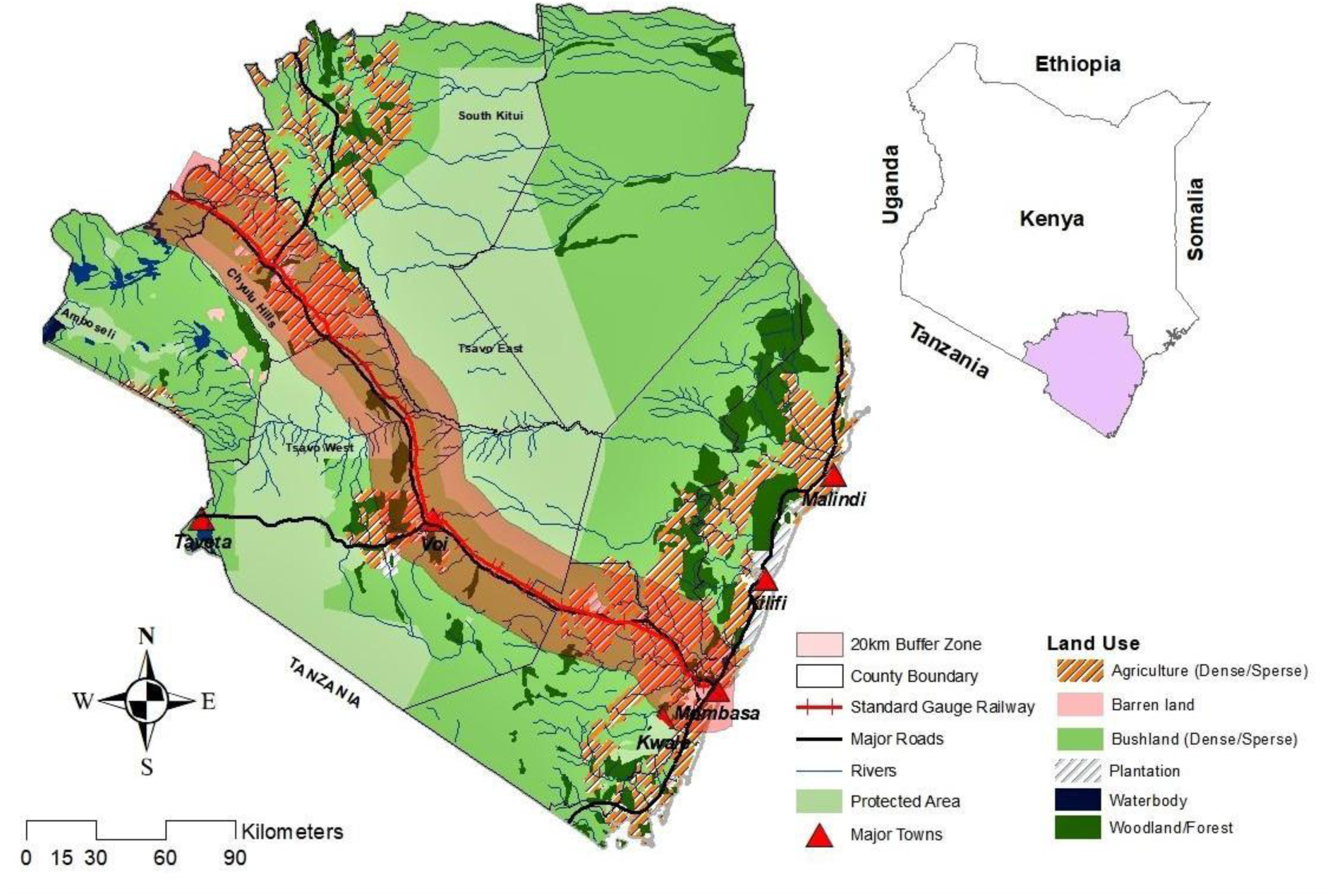
Map of the study area showing the 20 km buffer used for this study in southern Kenya.

The Tsavo landscape, on the other hand, comprises four National Parks and multiple community conservancies including Tsavo East and West National parks. Covering approximately 49,611 km^2^ Tsavo is the largest contiguous extent of savannah in Kenya. The ecosystem occupies parts of Taita Taveta and Makueni counties and is home to a wide variety of wildlife species including c. 40% of Kenya’s elephant population, 18% of Kenya’s black rhino population, and over 60 other mammal and 400 bird species (WRTI and KWS 2021) found in the protected areas and multiple community conservancies. The study landscape has a bimodal rainfall pattern with short rains from November to December and long rains in March, April and May. The average annual rainfall ranges between 200 and 700mm per annum. The average normal daily temperatures range between 20°C and 30°C.

The main source of livelihood for communities in the study area is agro-pastoralism. Locals mainly grow cassava, maize, sweet potato, pigeon peas and tree crops such as cashew nut and coconut while rearing cattle, goats, and sheep. Other livelihood activities in the area include charcoal production and informal sector businesses such as small retail shops and restaurants, and selling handicrafts, while others have joined formal employment locally or in other parts of Kenya (Kenya National Bureau of Statistics 2020). The study area is bisected by several linear infrastructure including the SGR, oil pipeline, power lines and highway, all of which run westwards from the coastal town of Mombasa through central Kenya to the Kenya-Uganda border in western Kenya (Nyumba *et al*. 2021). In addition, numerous feeder roads connecting to other regions in Kenya and south towards Tanzania criss-cross the study area. Although this study focused on major infrastructure within the study area, narrow scrutiny was placed on the recently built SGR, whose operation began in 2017. The SGR, unlike all other infrastructure in the study area, was designed with a wide range of safety protection measures that include speed limits, installation of a high guard fence, safety buffers and earth embankments to avoid crossing other infrastructures. In addition, bridges, underpasses, culverts and flyovers have been constructed in wildlife areas such as Tsavo and in high human density areas such as Mombasa to facilitate the free movement of wildlife and people (Nyumba *et al*. 2021). However, there have emerged claims that the SGR has become a barrier to wildlife movement deflecting wildlife to human settlements and increasing HWC.

### 2.2 Respondent sampling

Since we did not have a reliable household register from which we could draw our samples, we used spatial alongside proportional probability sampling techniques (Kondo *et al*. 2014, Pearson *et al*. 2015, Kassié *et al*. 2017, Chen *et al*. 2018), considering such factors as population density to represent human populations density rather than populations of space (e.g. Landry and Shen 2005). Gibson and Mckenzie (2007) detail the use of spatial data as a sampling tool in research. For this study, we developed a 20km buffer along the SGR, then subdivided the entire stretch based on administrative boundaries, in this case, counties and sub-location (Figure 1).

For each sub-location, we overlaid demographic data from the Kenya national census data for 2019 (KNBS 2019) to generate maps of population density and the number of households per sub-location. This enabled us to use the proportional probability sampling technique to determine the sample size for each sub-location within the target county. Using the “*Create Random tools*” in ArcGIS (Data Management tools>Feature class>Create Random Points), we generated random sampling points for each sub-location representing a household for the interview (ESRI 2011). **Figure 2(a)** is an example of the points generated for Taita Taveta County. A similar procedure was used for each of the counties within the study area. The digital maps were then uploaded onto mobile phones installed with the ‘Collector for ArcGIS’ App. The App enabled the enumerators to trace all the selected households and carry out the surveys. **Figure 2(b)** shows the actual location of selected households where actual questionnaires were administered on mobile devices held by the enumerators.

**Figure 2:**
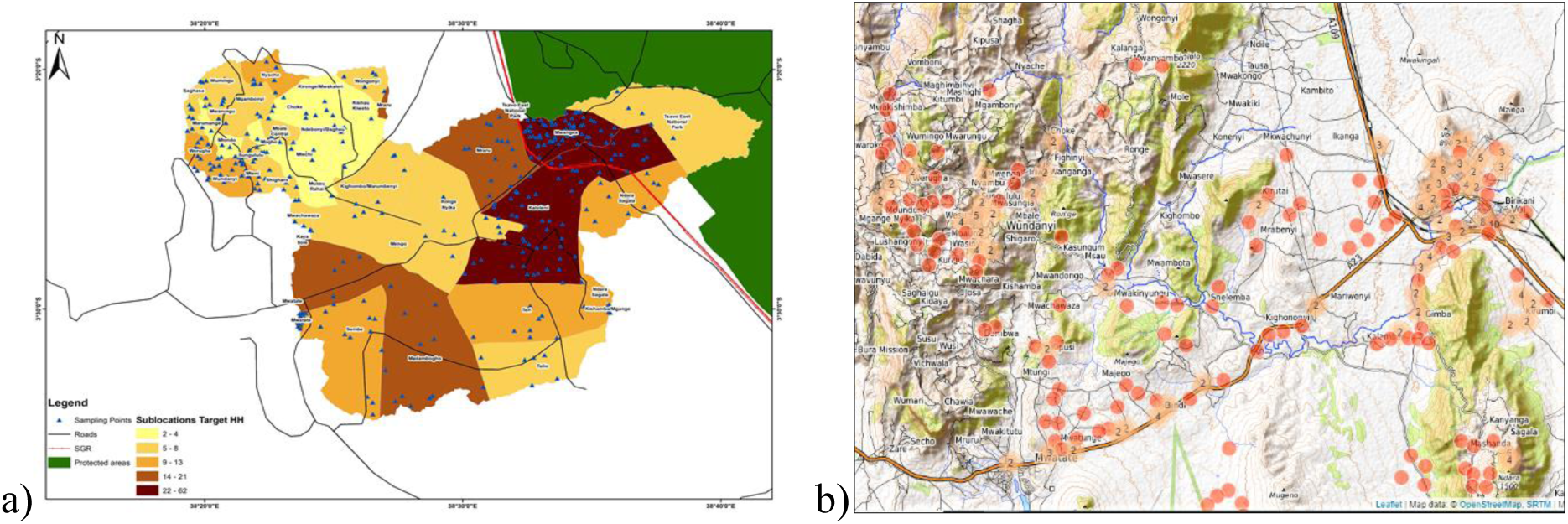
Random sampling points in Taita Taveta County generated through ArcGIS (a); and as seen on Collector for ArcGIS and Kobo Toolbox Web App (b).

### 2.3 Data collection

We designed our household data collection forms based on the Open Data Kit (ODK) suite. These included ODK build for form designer, ODK collect for user interface and Kobo toolbox for data storage services. We used MS Excel for the design of mobile forms, converted to XForms format using XLSFORM (https://xlsform.org) online tool, and validated using Enketo (https://enketo.org/xforms). We uploaded the validated form onto the Kobo toolbox server, a web-based application that supports mobile data collection and provides hosting services for data collected through mobile devices. We recruited and trained a team of field enumerators and provided them with android mobile phones installed with the ODK collect app. Each enumerator downloaded the digital forms for data collection from the Kobo toolbox server onto their devices for the survey. Completed survey forms were submitted to the server and our research teams tracked and verified the quality of the submitted data in real-time or near real-time and downloaded the data in comma-separated values (CSV) format for further analysis. We used the statistical program IBM SPSS Statistics for Windows, version 23 (IBM Corp 2013) for all statistical analyses and ArcGIS (ESRI 2011) for spatial analysis. We used generalized linear models (GLMs) in SPSS to explore a range of factors determining the probability of experiencing HWC using distance from the respondent’s home/property to different types of infrastructure as the baseline for comparison. The infrastructure under consideration were the SGR, oil pipeline, highway and power line. For this study, continuous explanatory variables included age and distance from the target infrastructure measured in kilometres (Km), whereas categorical explanatory variables included gender, education level, length of residency and main occupation.

### 2.4 Analysis

#### 2.4.1 Differences between administrative counties

We compiled boundaries of the counties of Kenya within ARC GIS. We then used spatial analysis tools to analyse if there was a significant difference in HWC between counties.

#### 2.4.2 Impact of distance to infrastructure

For purposes of this analysis, we defined distance from the nearest linear infrastructure using two criteria. First, we relied on respondents’ estimation of the distance from their property to the target linear infrastructure (SGR, oil pipeline, highway and power line); and then we used the proximity analysis tools in ArcGIS to calculate the actual distance from the property to different kinds of infrastructure, based on the GPS location of the property. Using these data we also generated heat maps to identify locations prone to HWC along the SGR. Furthermore, we modelled the influence of various linear infrastructure in the study area to establish their influence either singly or in combination with each other and other factors. Two models were created. The first considered distance from the SGR, oil pipeline, highway and power line separately. The second included respondent age, education, gender, occupation and length of residency.

#### 2.4.3 Correlation with other factors

We also used GIS and statistical approaches to analyse HWC incidents in relation to different land cover types (agricultural and plantation areas) and in relation to protected area boundaries.

## 3. Results

### 3.1 Respondent characteristics

Makueni County had the largest number of respondents (n=427, 35.0%) while Mombasa-Kwale-Kilifi and Taita Taveta counties had an almost equal number of respondents at n=395, 32.4% and n=398, 35.6%, respectively. Most of the respondents were females, although slightly more males were interviewed in Mombasa-Kwale-Kilifi counties. Most respondents were between 18 and 35 years old. Most respondents were either farmers or businesspersons.

### 3.2 Types of human-wildlife conflict between counties

Significantly (χ^2^=83.597, df=2, p<0.001) more respondents (n=881, 72% vs n=339, 28%) reported they had experienced HWC over the past year and these were grouped into six distinct types. Crop damage (40.28%) was the most prevalent conflict type, followed by livestock deaths (23.29%), livestock injuries (15.59%), human injuries (13.31%), and property damage (4.38%) and finally, human deaths (3.15%). Elephants and predators, especially lions, hyenas and wild dogs were the most prominent HWC species and hence the prominence of damage to crops and wildlife attacks on humans and livestock. However, there was a significant difference (*χ^2^=*238.467*, df=16, p*<0.001) in wildlife species involvement in HWC by counties (**Figure 3**).

**Figure 3:**
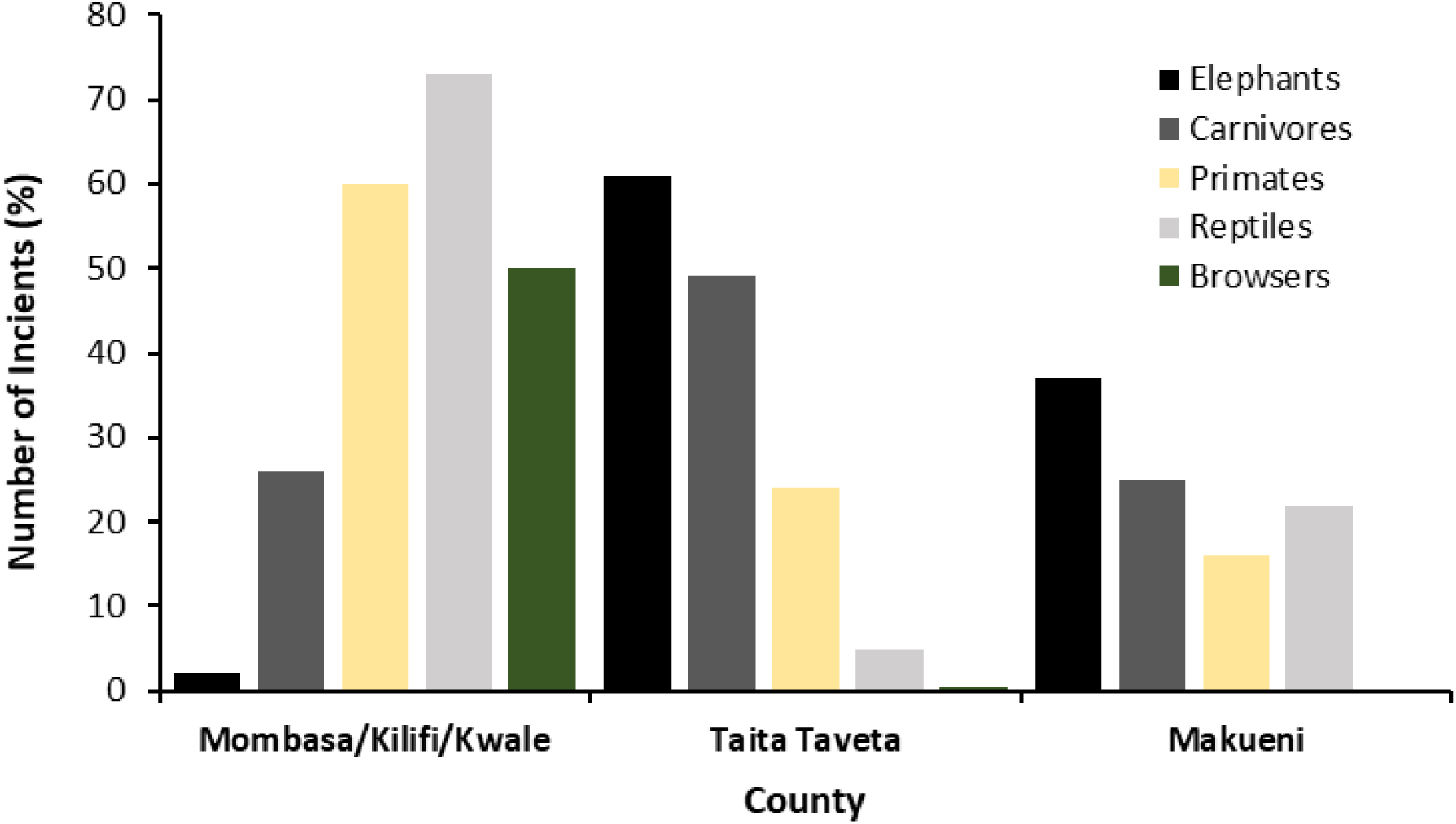
Human-wildlife conflict by type(s) of wildlife in each county.

### 3.3 Temporal patterns of human-wildlife conflict

Overall, most of our respondents felt that HWC had increased or stayed the same over the last five years. This is a critical period since the construction and operation of the SGR began during the five years up to the time of this survey. The change in HWC significantly varied (χ^2^=130.398, df=6, p<0.001) by county. Whereas Taita Taveta and Makueni counties reported a significant increase, HWC remained the same or increased marginally in Mombasa, Kwale and Kilifi counties. Between 15-25% of respondents could not tell whether there had been any changes in HWC (**Figure 4**).

**Figure 4:**
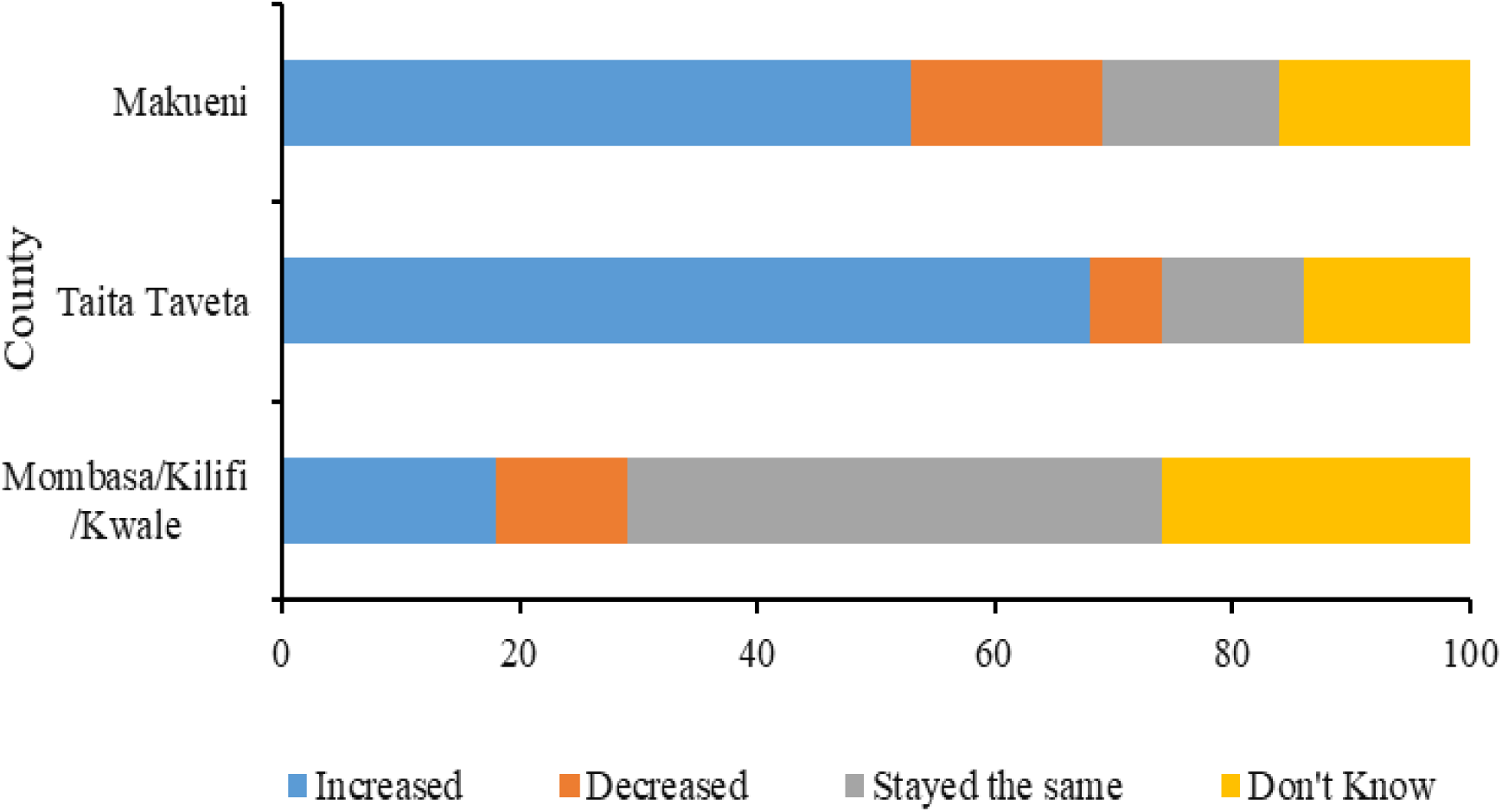
Change in human-wildlife conflict over the last five years by county.

### 3.4 Spatial patterns of human-wildlife conflict

Results showed a distinct relationship between HWC and distance from the target linear infrastructure. Respondents’ estimation ranged between 1.0-20.0 km with a mean of 5.3 km while proximity analysis ranged between 0.5-19.8km with a mean of 5.8km. Our results further indicated that the number of HWC incidents changed with distances from the target linear infrastructure to the property. Specifically, the number of incidents diminished as the distance from the SGR increased (57%) with a much less significant relationship of HWC increasing with increased distance from the oil pipeline (16%). Meanwhile, changes in distances from the highway and power lines did not result in any change in HWC (**Figure 5**).

**Figure 5:**
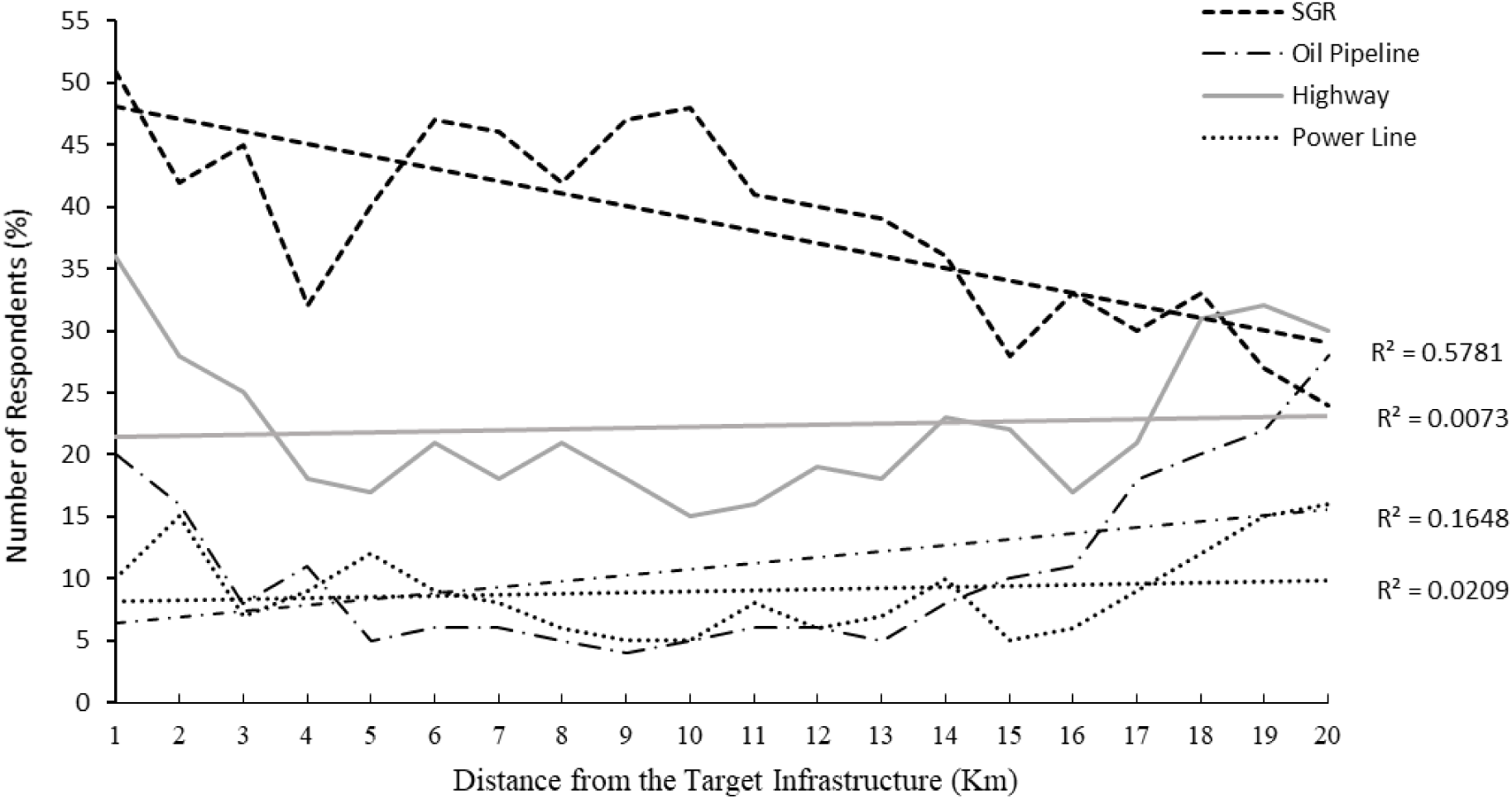
Number of households reporting HWC at different distances from the target infrastructure types.

Maps show a higher concentration of HWC closer to the SGR in all the study areas. The higher incidents of HWC between 16 – 20 km from the highway may be symptomatic of crop damages, livestock deaths, livestock injuries, or human injuries in agricultural and plantation areas and areas closer to the protected area boundary. **Figure 6** outlines the spatial clustering by species groups across the study area.

**Figure 6:**
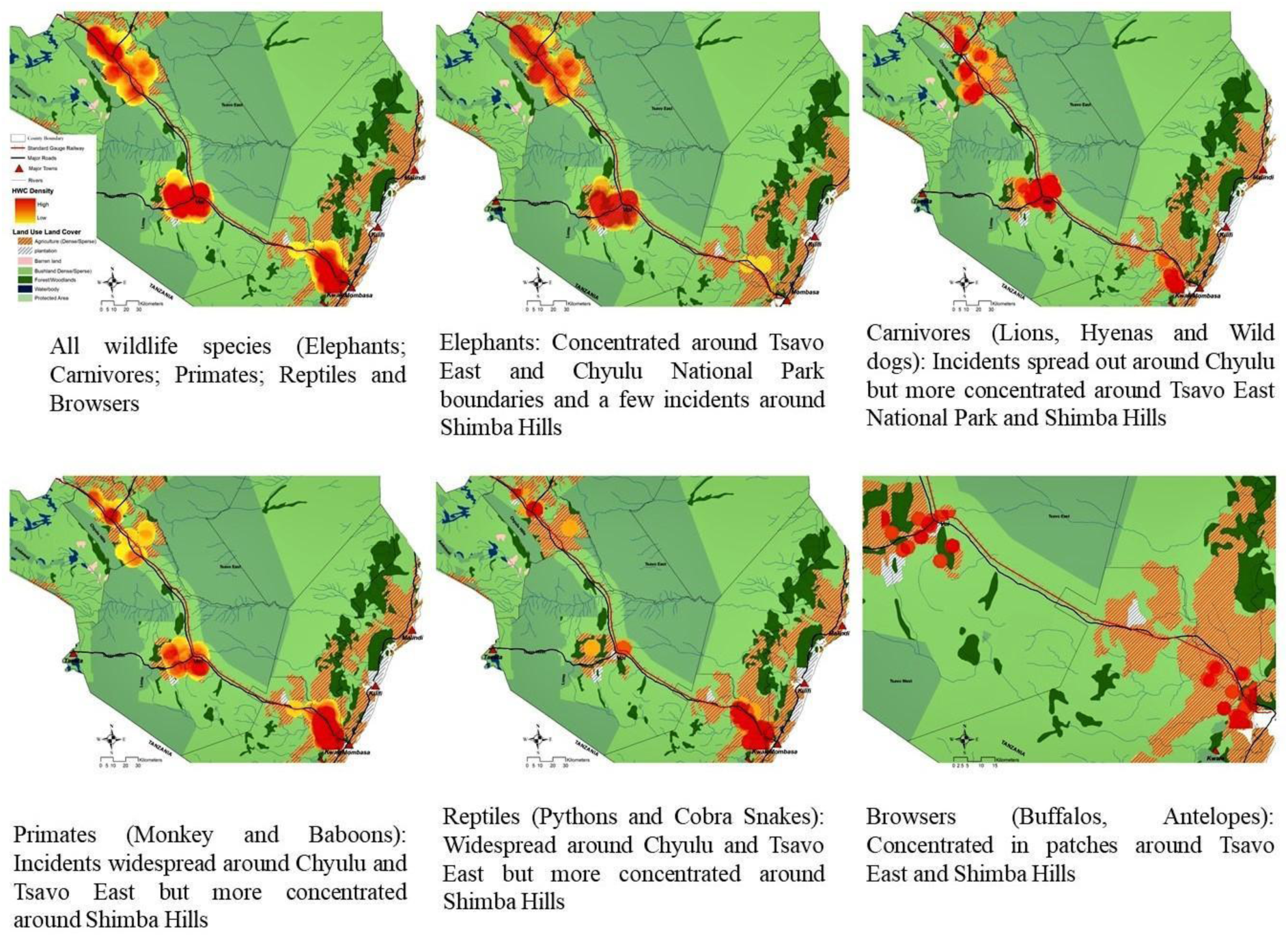
Heat map of human-wildlife conflict locations along the SGR and within different land use land covers.

### 3.5 Influence of distance from linear infrastructure on human-wildlife conflict

Our model was significant for all the types of infrastructure, suggesting that respondents’ level of education, age, gender, main occupation, length of residency and distance from respondents’ property to the target infrastructure significantly influenced the odds of experiencing HWC in the study area (**Table 1**). Our results in **Table 1** show that HWC decreased by 1% as you move away from the SGR and increased by 2% as you moved away from the oil pipeline. Highway and power lines did not influence HWC occurrence.

**Table 1:**
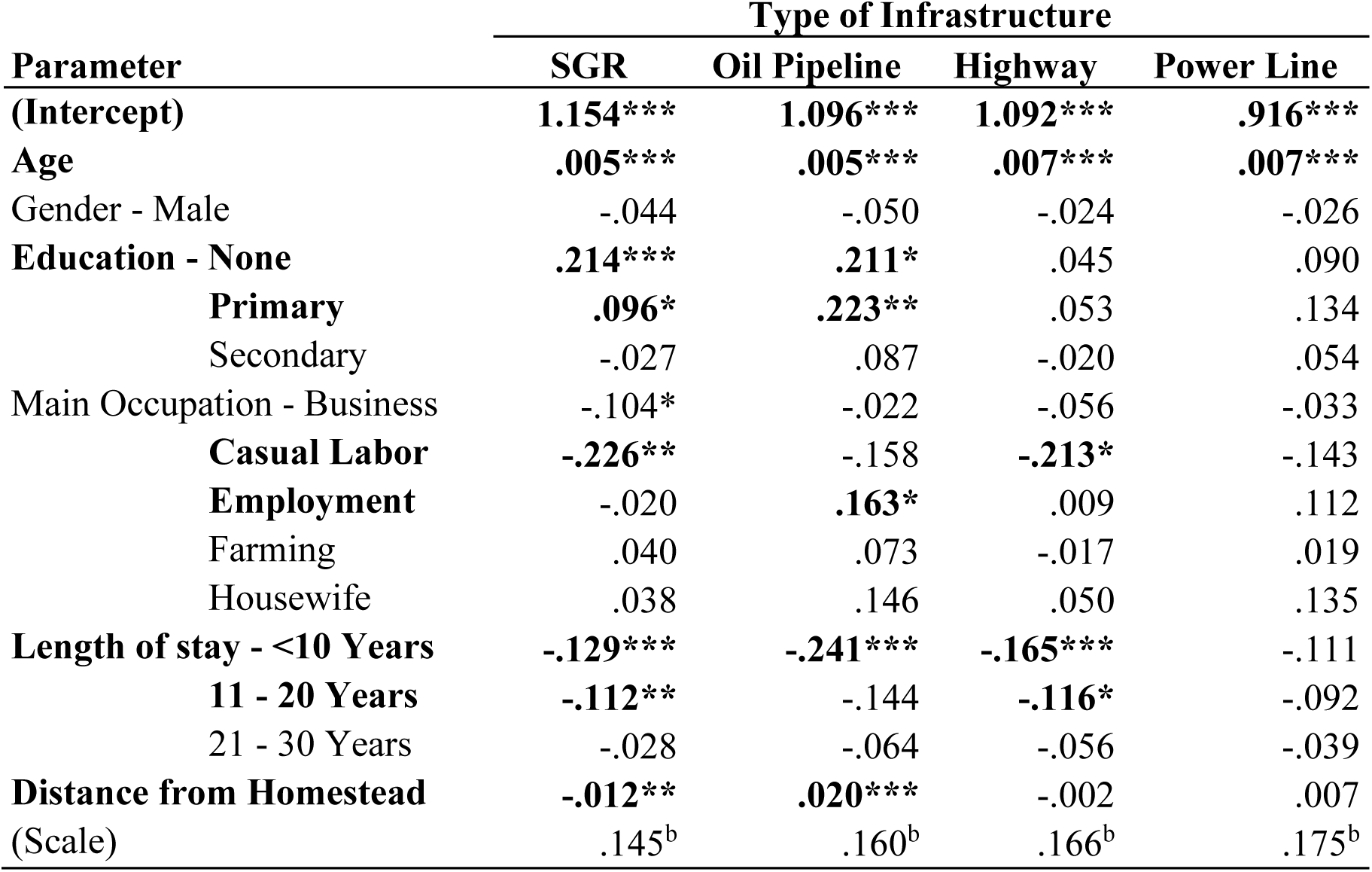
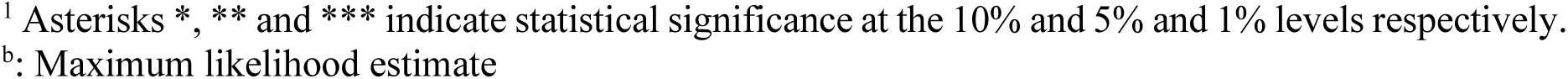
Generalized linear model results for factors influencing HWC under different types of infrastructure^1^.

| Parameter | Type of Infrastructure |  |  |  |
| --- | --- | --- | --- | --- |
|  | SGR | Oil Pipeline | Highway | Power Line |
| **(Intercept)** | **1.154\*\*\*** | **1.096\*\*\*** | **1.092\*\*\*** | **.916\*\*\*** |
| **Age** | **.005\*\*\*** | **.005\*\*\*** | **.007\*\*\*** | **.007\*\*\*** |
| Gender - Male | -.044 | -.050 | -.024 | -.026 |
| **Education - None** | **.214\*\*\*** | **.211\*** | .045 | .090 |
| **Primary** | **.096\*** | **.223\*\*** | .053 | .134 |
| Secondary | -.027 | .087 | -.020 | .054 |
| Main Occupation - Business | -.104* | -.022 | -.056 | -.033 |
| **Casual Labor** | **-.226\*\*** | -.158 | **-.213\*** | -.143 |
| **Employment** | -.020 | **.163\*** | .009 | .112 |
| Farming | .040 | .073 | -.017 | .019 |
| Housewife | .038 | .146 | .050 | .135 |
| **Length of stay - <10 Years** | **-.129\*\*\*** | **-.241\*\*\*** | **-.165\*\*\*** | -.111 |
| **11 - 20 Years** | **-.112\*\*** | -.144 | **-.116\*** | -.092 |
| 21 - 30 Years | -.028 | -.064 | -.056 | -.039 |
| **Distance from Homestead** | **-.012\*\*** | **.020\*\*\*** | -.002 | .007 |
| (Scale) | .145<sup>b</sup> | .160<sup>b</sup> | .166<sup>b</sup> | .175<sup>b</sup> |

When we considered all the types of infrastructure together, the influence of different factors changed drastically. Whereas having no education or primary level of education influenced HWC when different types of infrastructure were analysed separately at 21% and 10%, respectively, attaining primary and secondary levels of education became more influential when different types of infrastructure were analysed together at 27% and 21%, respectively (**Table 2**). The plausible reason could be the increased reliance of humans and their livelihood systems on natural resources, thus, more interactions with wildlife among those with low or no formal education. More importantly, the results showed that the influence of the SGR diminished when different types of infrastructure were analysed together. **Table 2** accounts for the changes in the influence of different factors when different types of infrastructure were analysed together.

**Table 2:** Generalized linear model results for factors influencing HWC occurrence based on distances from the nearest linear infrastructure.

| Parameter | B | Std. Error | Hypothesis Test |  |  |
| --- | --- | --- | --- | --- | --- |
|  |  |  | Wald Chi-Square | df | Sig. |
| **(Intercept)** | **1.059** | **.1851** | **32.731** | **1** | **.000** |
| **Age** | **.006** | **.0024** | **7.105** | **1** | **.008** |
| Gender - Male | -.112 | .0692 | 2.613 | 1 | .106 |
| Education - None | .098 | .1348 | .528 | 1 | .467 |
| **Primary** | **.265** | **.1057** | **6.268** | **1** | **.012** |
| **Secondary** | **.207** | **.1030** | **4.025** | **1** | **.045** |
| Main Occupation - Business | -.118 | .1107 | 1.135 | 1 | .287 |
| **Casual Labour** | **-.422** | **.1748** | **5.826** | **1** | **.016** |
| Employment | .173 | .1267 | 1.859 | 1 | .173 |
| Farming | -.015 | .0979 | .022 | 1 | .881 |
| Housewife | .053 | .1261 | .176 | 1 | .675 |
| **Length of stay - <10 Years** | **-.318** | **.1032** | **9.479** | **1** | **.002** |
| **11 - 20 Years** | **-.247** | **.0923** | **7.148** | **1** | **.008** |
| **21 - 30 Years** | **-.152** | **.0896** | **2.888** | **1** | **.089** |
| Distance (Km) - SGR | .008 | .0156 | .263 | 1 | .608 |
| Oil Pipeline | -.010 | .0139 | .512 | 1 | .474 |
| Highway | -.016 | .0130 | 1.609 | 1 | .205 |
| **Power Line** | **.021** | **.0126** | **2.783** | **1** | **.095** |
| (Scale) | .136<sup>b</sup> | .0151 |  |  |  |

<sup>b</sup> Maximum likelihood estimate.

## 4. Discussion

HWC research has been undertaken over the past four decades (Naughton-Treves 1998, Hill 2004, Treves 2006) and new information continues to emerge that merits intense study to enhance a better understanding of HWC. One of the emerging and fast-growing drivers of HWC is the growth and expansion of linear transportation infrastructure that is increasingly transforming wildlife habitats worldwide. Currently, however, limited literature exists on the influence of linear transportation infrastructure on HWC, as most studies have addressed linear infrastructure within the broader context of ecological impacts on wildlife and not human-wildlife interactions (Anderson, 2002; Forman et al., 2003). However, negative interactions between humans and wildlife continue to pose challenges to co-existence in multifunctional landscapes like Tsavo and Shimba Hills in Kenya.

Several previous studies within our study area have identified damage to crops and attacks on livestock as the most important HWC (Ngure 1992, Sindiga 1995, Mwamidi *et al*. 2012, Long *et al*. 2020, Lala *et al*. 2021). Human activities in the area are dominated by crop farming and pastoralism, production activities that conflict with wildlife conservation across wildlife ranges (Hoft and Hoft 1995, Graham *et al*. 2010, Kariuki *et al*. 2021). Coupled with recent growth and expansion of linear infrastructure, incidents of HWC seem to be on the rise (e.g. Long *et al*. 2020). Munyao *et al*. (2020) ranked general threats, especially from elephants as the most important type of HWC in Tsavo. Threats from wildlife derive from the presence of wildlife within human settlements that do not result in any of the other types of HWC but might include imposing fear on people through aggressive behaviours. Such presence could be linked to wildlife moving into the human settlement to avoid the busy linear infrastructure networks across the landscapes. Okita-Ouma *et al*. (2017) and Nyumba *et al*. (2021) established that elephants in Tsavo had become aggressive, because of disturbance from the increased use of the linear transport infrastructure across the landscape.

HWC tends to follow distinct spatial and temporal patterns usually linked to the time of day, seasons, proximity to wildlife refuges and other key resources (Ackers *et al*. 2001, Gupta 2011, Prasad and Hiny 2011, Long *et al*. 2020, Siljander *et al*. 2020). However, our study demonstrated that distance from linear transportation infrastructure including railways, highways, oil pipelines and power lines to human settlements and property could also affect HWC patterns. We showed that HWC diminished with distance from the railway line, while the distance from the highway, oil pipeline and electricity lines had little influence. The railway, unlike the highway, is a recent development project conceived, planned and constructed at a higher political premium (Government of the Republic of Kenya and the National Economic and Social Council 2007) and hence publicity in Kenya, is mostly negative and controversial (Kamau 2015, Rajab 2017, Wafula 2018, Nyumba 2021). Since our data was based on human perceptions and assessment, and hence constructed meanings (Knight 2000), the assessment of HWC patterns could have been influenced by their understandings, values and agendas regarding wildlife conservation and development in the landscape.

The spatial patterns of HWC in the study area were aligned with the railway line and the highway. Indeed, our data showed that HWC incidents were clustered within 5.3km of the railway and within areas of intense farming and plantation agriculture (Figure 6). This observation is important but must be discussed against the background of traditional land use land cover systems and historical changes in HWC in the landscape. A recent land-use land cover change analysis along the entire SGR corridor established that cropland and the built-up areas had increased between 2010-2019, a period that coincided with the construction of the SGR (Sang *et al*. 2022). Nyumba *et al*. (2021) found that human settlements and associated activities like farming and livestock keeping had encroached on the way leaves and road reserves bisecting the Tsavo National parks along the SGR corridor. Coupled with the technical design of the SGR which created permanent embankments causing substantial hindrance to wildlife movements (Okita-Ouma, Lala, Koskei, *et al*. 2017, Munyao *et al*. 2020, Nyumba *et al*. 2021), it is intuitive to expect that wildlife, including conflict species, will be attracted to and confined within these locations and hence incidents of HWC.

We must also consider the species-focused influence of the SGR and highways on HWC. The results of this study indicate that different conflict species depicted different cluster patterns in different locations. Long-term wildlife census and wildlife movement data from the Tsavo and Shimba landscapes show different species’ responses to infrastructure, especially elephants and other large mammals (Okita-Ouma *et al*. 2016, Okita-Ouma, Lala, Koskei, *et al*. 2017, Okita-Ouma, Lala, Moller, *et al*. 2017, Munyao *et al*. 2020, Lala *et al*. 2022). The studies observed that wildlife movement across the SGR was curtailed and with most of them wandering into human settlements while finding their new path to safe refuges. From our data, species that are sensitive to road traffic such as primates and browsers appeared to cluster away from the railway and highway (Figure 6) while bold conflict species like elephants and carnivores clustered closer to the railway and highways. This finding does not imply that the SGR and highway determined the occurrence and distribution of HWC but could lend themselves to further analysis of the influence of infrastructure both in isolation and in the presence of other drivers.

Historically, socio-demographic drivers of HWC such as age, gender, occupation, residency, and education have been studied and remedies sought (e.g. Hoare 2015). The present study shows that linear transportation infrastructure could be one of the drivers of HWC in multi-functional landscapes. Our data shows that HWC is influenced by distance from the SGR and oil pipeline, such that the number of incidents diminished by 1% as you move away from the SGR. This observation aligns with the observed wildlife, especially elephant, distribution in response to threats of poaching. Studies have shown that elephants coalesced around roads in Tsavo and Marsabit where the poaching threat was less than in vegetated areas where poachers enjoyed more cover (Ngene *et al*. 2009, 2011). This narrows the distances and, in a landscape, where farming activities are also concentrated near the roads and railways, incidents of HWC then become unavoidable.

It might be expected that existing policy mechanisms, such as the environmental impact assessments (EIA) or strategic environmental assessments (SEA), should identify and mitigate the effects of LTIs. However, EIAs often place the burden of proof on project opponents with insufficient ecological information and knowledge to determine the actual effects of LTIs, are limited in scope on the direct effects of LTIs only and are based on local considerations without broader landscape-level context (Gullett 1998, Retief *et al*. 2020). The Kenya National Human-Wildlife Coexistence Strategy and Action Plan 2024-2033 places priority on land and space management, innovative mitigation strategies, capacity development, reforming institutions to improve coordination, and compensation for losses incurred due to human-wildlife conflicts. While this may be a good fodder for human-wildlife co-existence, it’s more urgent to ensure a robust framework to mitigation risks from human-wildlife conflicts through avoiding or minimising their occurrence and offsetting loss of biodiversity, particularly those from infrastructure barriers.

Strategic environmental assessments do have greater potential to capture the wider impacts and how these are connected to existing communities and land uses at a landscape level. Clearly, the development of linear infrastructure does lead to increased HWC and there are spatial and animal / community specific patterns to this conflict. Such increased insight into the unintended HWC consequences of transport and linear infrastructure development can be used to mitigate these and maximise the social, economic benefits while not compromising wildlife and natures contributions to people.

## 5. Conclusion

Conflicts between humans and wildlife continue to threaten nature conservation and human development globally. HWC severely threaten the food security and livelihoods of smallholder households sharing space with or living adjacent to wildlife conservation areas such as the Tsavo and Shimba Hills landscapes in southern Kenya. These conflicts are driven by several ecological and anthropogenic factors acting singly or in combination. Recently, the rapid growth and expansion of infrastructure have added another layer of drivers, especially in multifunctional landscapes such as Tsavo and Shimba Hills. Within these landscapes, we identified six types of conflicts, but attacks on livestock and crops remain the most important for residents. The number of HWC incidents diminished with distance from the SGR line such that the closer a property is to the line, the more vulnerable it is to HWC. This, however, varied by species, such that elephants and predators attacked property closer to the railway line compared to primates and other browsers as determined by the spatial patterns of HWC. Nonetheless, the influence of the SGR on HWC is very minimal and this is even further obscured when considered in the presence of other known factors such as education, gender, length of residency, age and occupation.

The use of people’s perceptions and assessment of HWC based on recall and personal and communal experiences is a well-recognised research technique. Our use of this technique was driven by the fact that, to our knowledge, few or no systematic studies have explored the linkages between HWC and linear transportation infrastructure, especially in sub-Saharan Africa. But we believe that this study provides some foundational data and information that can inform the need for a more long-term, landscape-level study to explore deeply the influence of LTI. We propose that such a study should be based on field-based scientific monitoring of HWC incident, telemetry data and wildlife movement patterns and trends over time and systematically linking them to the existence and use of the LTI, incorporating more ecological determinants, species behavioural factors and biophysical factors such as elevation and climate variations, to filter out the specific role of LTI in HWC.

## 6. Acknowledgement

First and foremost we acknowledge the outstanding contribution of Dr. Tobias Nyumba to research on the ecological impacts of large-scale infrastructure development in Africa. Following his PhD at the University of Cambridge, Tobias held a Postdoctoral Fellowship with the Development Corridors Partnership at the African Conservation Centre and the University of Nairobi, where his work focused on transportation infrastructure and its implications for biodiversity where much of the data for the current manuscript were generated. We acknowledge funding from the UK Research and Innovation’s Global Challenges Research Fund (UKRI GCRF) through the Development Corridors Partnership project (Project No. ES/P011500/1). Subsequently, as a Marie Skłodowska-Curie Fellow at the University of York, Tobias advanced participatory scenario planning approaches, including the application of the ‘KESHO’ tool, to explore alternative land-use futures. Through training workshops and extensive stakeholder engagement, Tobias successfully bridged science and policy, influencing conservation practice in Kenya and beyond. Tobias’s scholarly contributions, passion and commitment to ensuring space for wildlife in dynamic socio-economic contexts continue to inform conservation science and practice.

